# Modeling metastatic progression from cross-sectional cancer genomics data

**DOI:** 10.1101/2024.01.30.577989

**Authors:** Kevin Rupp, Andreas Lösch, Y. Linda Hu, Chenxi Nie, Rudolf Schill, Maren Klever, Simon Pfahler, Lars Grasedyck, Tilo Wettig, Niko Beerenwinkel, Rainer Spang

**Author notes:** These authors contributed equally (Kevin Rupp, Andreas Lösch, Y. Linda Hu).

## Abstract

Metastasis formation is a hallmark of cancer lethality. Yet, metastases are generally unobservable during their early stages of dissemination and spread to distant organs. Genomic datasets of matched primary tumors and metastases may offer insights into the underpinnings and the dynamics of metastasis formation. We present metMHN, a cancer progression model designed to deduce the joint progression of primary tumors and metastases using cross-sectional cancer genomics data. The model elucidates the statistical dependencies among genomic events, the formation of metastasis, and the clinical emergence of both primary tumors and their metastatic counterparts. metMHN enables the chronological reconstruction of mutational sequences and facilitates estimation of the timing of metastatic seeding. In a study of nearly 5000 lung adenocarcinomas, metMHN pinpointed TP53 and EGFR as mediators of metastasis formation. Furthermore, the study revealed that post-seeding adaptation is predominantly influenced by frequent copy number alterations. All datasets and code are available on GitHub at https://github.com/cbg-ethz/metMHN.

## 1 Introduction

Metastasis is the primary cause of cancer-related death. It occurs as tumors evolve, when the primary lesion extends beyond its initial boundaries, invading adjacent healthy tissues, lymph nodes, and blood vessels. Cancer cells can then enter the bloodstream and spread to different locations within the body. At these new sites, the disseminated cells face novel selective pressures, leading to the elimination of many, but not all, cells. The survivors adapt and eventually colonize these foreign tissues, forming metastases [23].The development of cancer, or tumorigenesis, is predominantly driven by the progressive accumulation of genomic alterations, including somatic mutations and copy number alterations in cancer driver genes [40]. These alterations often result in divergent genotypes between a primary tumor and its associated metastasis. Extensive clinical sequencing efforts like the MSK-MET study [26] recently compiled genomic data from primary tumors and metastases. In principle, such datasets may inform about the timing and genetic mechanisms of metastasis formation, but revealing these pieces of information is challenging.

Cancer progression models aim to infer interactions between genomic alterations based on their co-occurrence patterns in cross-sectional data. Such models can then be used to both predict the future progression of tumors as well as to explain the past by inferring the order in which observed alterations accumulated. These models have their roots in the pioneering work of Fearon and Vogelstein [14]. Since then, a variety of models and algorithms have emerged to refine and expand upon this concept. They include Conjunctive Bayesian Networks [2], CAPRI [31], Network Aberration Models [19], HyperTraPS [18] and Mutual Hazard Networks [33]. All of these models only consider the progression of a single sequence and thus can not capture the divergent, branching patterns characteristic of metastatic disease progression. Therefore none of the above mentioned models can leverage the information provided by matched primary tumor and metastasis samples from the same patient. Methods like REVOLVER [7] or TreeMHN [24] can account for this branching behaviour as they model evolution of tumors on a clonal level. However, they require phylogenetic data and are not explicitly designed to model metastatic branching.

Here, we present Mutual Hazard Networks for metastatic disease (metMHN), a cancer progression model that captures the branching progression observed in primary tumors and their metastatic offshoots. The model is designed to infer interactions among genomic alterations and to assess their impact on the propensity for a tumor to seed a metastasis. Additionally, it accounts for metastasis-specific effects on the rates at which genomic alterations accumulate. metMHN utilizes both cross-sectional data from matched primary tumors and metastases, and singular observations of only one of the two. It also models how genomic changes affect tumor observability. We demonstrate the utility and robustness of the metMHN model using the lung adenocarcinoma dataset (LUAD) provided by the provided by the Memorial Sloan-Kettering Cancer Center through AACR GENIE [30].

## 2 Methods

metMHN extends the Mutual Hazard Network (MHN) framework, originally introduced by Schill et al. in 2020 [33] and further developed by Schill et al. in 2023 [34], which models the progression of primary tumors. We first establish the notation employed by MHNs and then introduce metMHN.

### 2.1 Mutual Hazard Networks

MHNs [33] model the evolution of primary tumors as a continuous-time Markov chain (CTMC) *{X*(*t*), *t ≥* 0*}* on the binary state space *{*0, 1*}*^*n*^. A state *x ∈ {*0, 1*}*^*n*^ corresponds to a set of progression events, such as mutations or copy number alterations, where *x*_*i*_ = 1 indicates that event *i ∈ {*1, …, *n}* is present, whereas *x*_*i*_ = 0 indicates its absence. Let 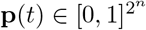 denote the probability distribution over states at time *t*, where the states are ordered lexicographically. The evolution of the probability distribution over time is governed by the Kolmogorov forward equation

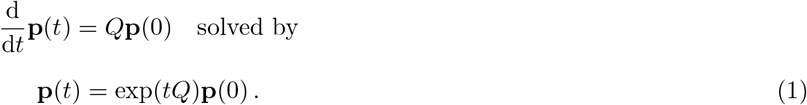

Here **p**(0) denotes the distribution over states at the start of the progression. It is assumed that all tumors start in a wild type state, where no event has occurred yet, thus 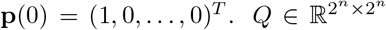 denotes the transition rate matrix on the state space. Events are assumed to accumulate irreversibly and one at a time. Therefore, the only non-zero off-diagonal entries of Q are the transition rates from states *x* = (…, *x*_*i−*1_, 0, *x*_*i*+1_, …) to *x*_+*i*_ = (…, *x*_*i−*1_, 1, *x*_*i*+1_, …) that differ by exactly one event *i*. The transition rates are parameterized by a much smaller matrix 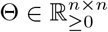 as

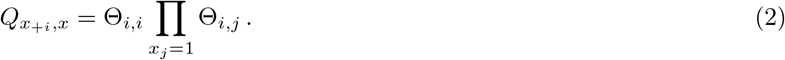

Here Θ_*i,i*_ denotes the base rate with which event *i* spontaneously occurs in a tumor and Θ_*i,j*_ the multiplicative effect of the presence of event *j* on the rate of event *i*. The age of a tumor at the time of its diagnosis is unknown. In [33] it is assumed to be exponentially distributed with mean 1 and independent of the state of the tumor. Marginalizing over *t* in Equation (1) yields the time-marginal distribution

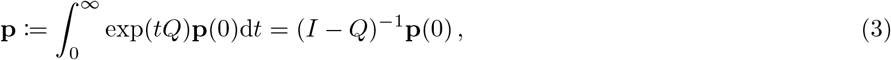

where *I* denotes the identity matrix. Let **p**_*x*_ denote the probability of observing a tumor in state *x*. Then the average log-likelihood for a dataset *𝒟* of tumor states is defined as

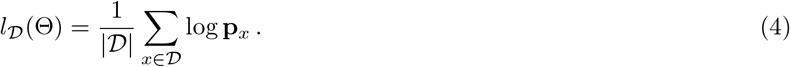

The matrix *Q* does not need to be stored explicitly, because it can be written as a sum of tensor products. By using tensor operations, **p** can be calculated efficiently and Θ can be learned with a time and space complexity only exponential in the number of events that have occurred for each tumor in the dataset, rather than exponential in 2*n* [32, 5]. Recently [22, 16, 28] reduced the complexity further to *n*^3^ using modern tensor formats and thus made MHN applicable to even larger state spaces.

Clearly, a tumor can only appear in a dataset after it has been clinically detected. This detection, in turn, is influenced by the tumor’s genotype, as certain mutations can induce growth or alter the tumor’s morphology. Such changes may result in symptoms that lead to the tumor’s discovery, followed by its diagnosis, surgical removal, and eventual sequencing. Therefore the rate of observation should be dependent on the state of the tumor. In [34], the observation of a tumor was introduced as a separate event with its own set of parameters 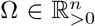. The observation of a state *x* happens at a rate 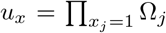, where Ω_*j*_ is a multiplicative effect of the presence of event *j* on the rate of observation. On the other hand multiplicative effects of the observation on other events are set to 0. Thus, as soon as the observation event occurs, progression is halted. States where the observation occurred are thus absorbing states of the Markov chain. Then the probability distribution at observation is equal to the stationary distribution **p**(*∞*) and given by

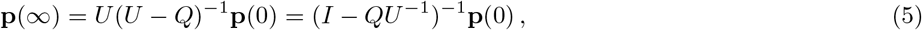

with 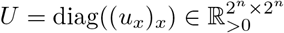 and *Q* and **p**(0) defined as in Equation (3) [34].

### 2.2 metMHN

We now present metMHN, an extension of the original MHN framework, meticulously tailored to analyze the dynamics of metastatic cancers.

#### 2.2.1 Dynamics on the combined state space

With metMHN, we model the joint progression of primary tumors and metastases as a Markov process on the combined event space of both tumor entities (see Figure 1b). Formally, we consider a CTMC *{X*(*t*), *t ≥* 0*}* on the state space *𝒮* := *{{*0, 1*} × {*0, 1*}}*^*n*^ *× {*0, 1*}*. A state *x ∈ 𝒮* is represented by a bit string of length 2*n* + 1. Each of the *n* progression events is encoded by two bits. The first bit 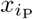 indicates the status of event *i ∈ {*1, …, *n}* in the primary tumor, and the second bit 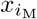 indicates the status of event *i* in the metastasis. We use the notations 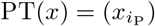 and 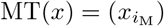 for *i* in *{*1, …, *n}* to refer to the genotypes of the primary tumor and the metastasis respectively. The (*n* + 1)^th^ event is encoded by one bit only and indicates the status of the seeding event. In the model context, the seeding event denotes that the progression of the metastasis has become decoupled from the progression of the primary tumor. Analogous to MHN we parameterize all transition rates by a low-dimensional set of parameters Θ *∈* ℝ^(*n*+1)*×*(*n*+1)^, where Θ_*i,i*_ refers to the base rate of event *i* and Θ_*i,j*_ to the effect of event *j* on the rate of event *i*. Before and after the seeding of a metastasis we assume different transition dynamics, which we describe in the following paragraphs.

**Figure 1:**
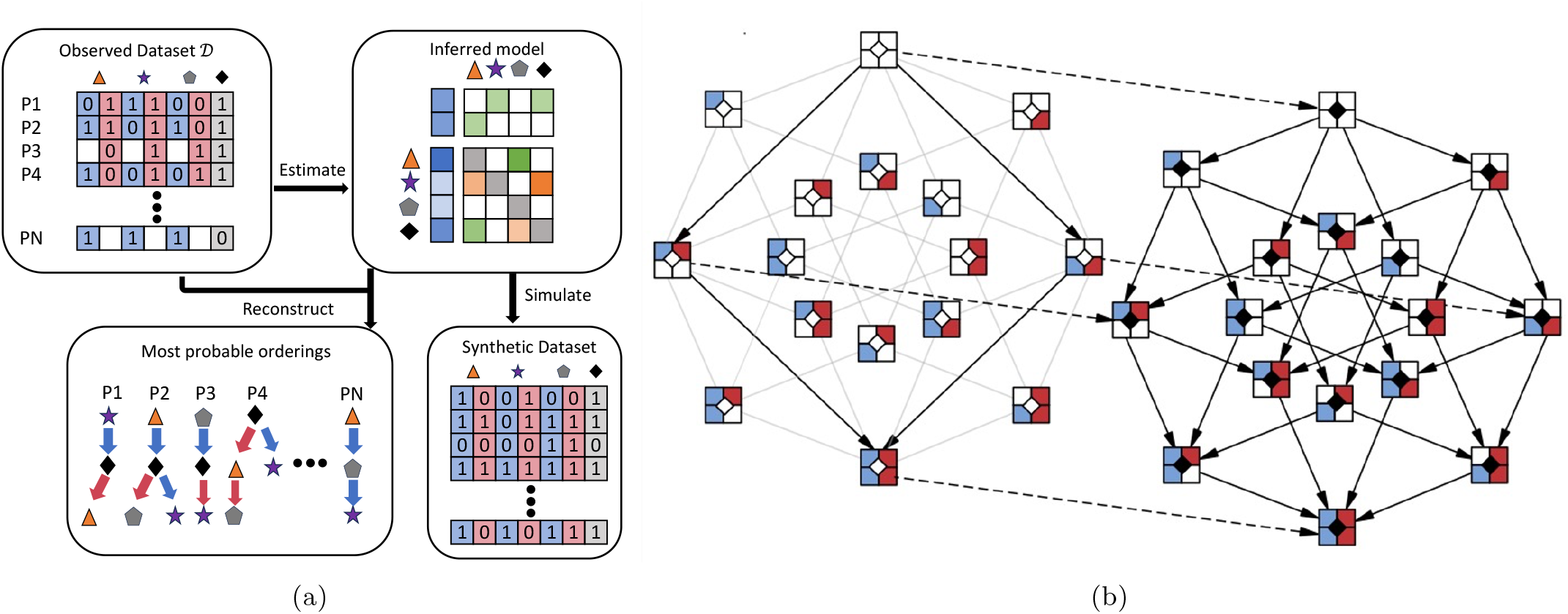
(a) Workflow of metMHN. In the top-left section, we show the types of input data that metMHN processes. Each row corresponds to a patient, each column to an event in the primary tumor (blue) or the metastasis (red). Events are represented by symbols and their status is encoded with a ‘1’ for present, ‘0’ for absent, or left blank if a tumor is unobserved. On the right, we present the primary output of metMHN: A network of interactions between events in matrix form. In the lower section, we show the most probable chronological ordering in which events accumulated in observed data points as inferred by metMHN. The progression trajectory of the primary tumor is indicated by blue arrows, while the trajectory of the metastasis is marked by red arrows. (b) The metMHN process and its state space: Black-bordered squares represent full states: the two compartments on the left detail the status of the primary tumor, the two on the right correspond to the metastasis, and the central diamond symbolizes the seeding event. The diagram is divided into two subspaces, with the left half constituting the subspace *𝒮*_0_ and the right half comprising the subspace *𝒮*_1_. Transitions between states that occur at non-zero rates are shown as solid black arrows. Transitions that are not possible in *𝒮*_0_ but are possible in *𝒮*_1_ are indicated by greyed-out arrows. Dotted arrows highlight transitions that influence the seeding event specifically.

Prior to seeding, the (soon-to-be) metastasis is identical to the primary tumor. Thus, events occur simultaneously in the primary tumor and the metastasis. Formally, we can describe these dynamics by a CTMC on the subspace *𝒮*_*0*_ := *{{*0, 1*} ×{*0, 1*}}*^*n*^ *×{*0*} ⊂ 𝒮* with transition rate matrix 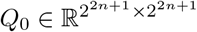. Let 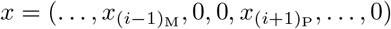 and 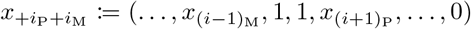 be states that differ by exactly one event *i*. Transitions from states *x* to states 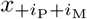 happen at rate

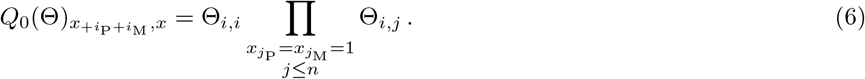

All other transitions within *𝒮*_0_ are prohibited (rate 0).

After seeding, the primary tumor and the metastasis are separate tumors and we assume that both accumulate mutations independently of each other. Formally, we describe the post-seeding dynamics by a CTMC on the sub-space *𝒮*_1_ = *{{*0, 1*} × {*0, 1*}}*^*n*^ *× {*1*} ⊂ 𝒮*. We introduce two transition rate matrices *Q*_P_ and 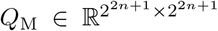.*Q*_P_ holds the rates for transitions that change only the primary tumor part of a state *x*: Transitions from states 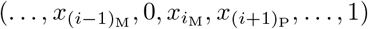 to states 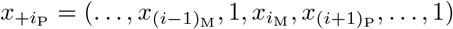occur at rate

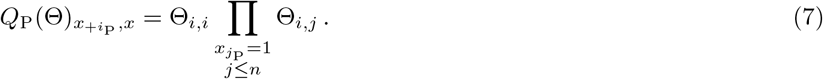

Note that transition rates in *Q*_P_ only depend on the primary tumor genotype PT(*x*) and not on the full state *x*. Since events must occur one at a time, all other transitions on *𝒮*_1_ that affect the primary tumor occur at rate 0. *Q*_M_ holds the rates for transitions that change only the metastasis part of a state *x*. We assume that metastatic tumors spread to foreign sites and face novel selective pressures that can differ drastically from the original site. We account for this by explicitly modeling effects from the seeding event on the progression events. Progression events occur in the metastasis at a rate given by the product of their base rates, the effects of events that are present in the metastasis and the effect of the new environment. Hence, transitions from states 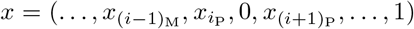 to states 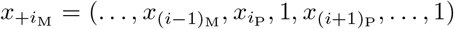 occur at rate

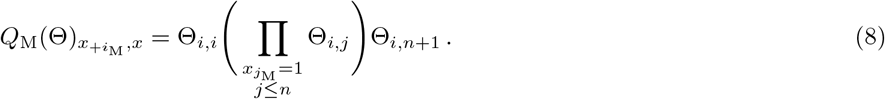

All other transitions on *𝒮*_1_ that affect the metastasis are prohibited (rate 0). The full transition rate matrix on *𝒮*_1_ is then given by the sum of *Q*_P_ and *Q*_M_.

By construction, the last event that occurs jointly and at the same time in a primary tumor and metastasis is the seeding event. Let 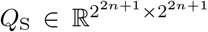 denote the transition rate matrix that holds the rates for all transitions from states 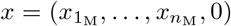, 0) in *𝒮*_0_ to their corresponding states 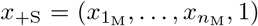 in *𝒮*_1_. Such transitions occur at rate

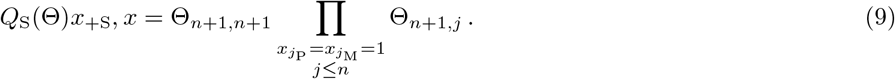

See Figure 1b for an illustration of the state space for *n* = 2. The transition rate matrix on the full state space *𝒮* is then

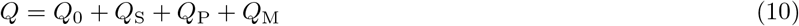

and we denote the probability distribution over states at time *t* by **p**(*t*). Following [22] we also provide formulas for the matrices *Q*_0_, *Q*_S_, *Q*_P_, *Q*_M_ as sums of tensor products in the supplement. By using these tensor structures in conjunction with the methods outlined in [32], the model parameters can be learned with a time and space complexity only exponential in the number of events that have occurred for each sample in the dataset, rather than exponential in 2(2*n* + 1).

#### 2.2.2 Modeling consecutive observations

Following [34] we model the observation of tumors explicitly as events. Since we model two tumors that at some point evolve independently and can also be observed separately, we have to include two distinct observation events. Thus we now model a CTMC on the extended state space *𝒮*_*D*_ := *𝒮 × {*0, 1*}*^2^. Let 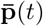 denote the probability distribution over states on the extended state space at time *t*. We assume that each event has a multiplicative effect on the rate of observation of the tumor it occurred in. Since the events that lead to the detection of a primary tumor can be vastly different from the effects that lead to the detection of a metastasis, we introduce two separate parameter vectors 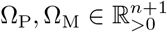 that contain the effects of progression events in the primary tumor and the metastasis on the rates of their respective observation event.

The primary tumor and the metastasis observation rates are defined as

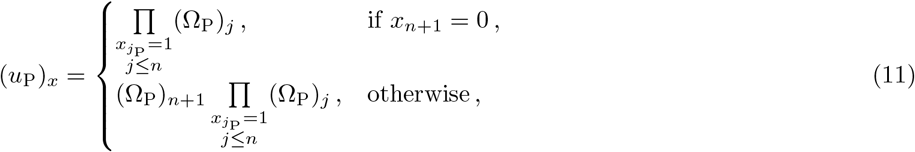

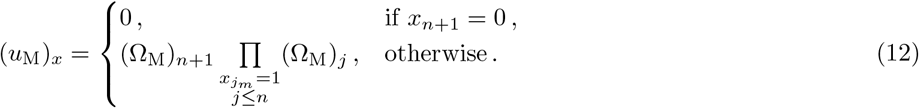

Let 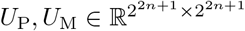 denote the diagonal matrices that hold the observation rates for primary tumors and metastases respectively and *U*_S_ = *U*_P_ + *U*_M_. We define that a metastasis is not observable prior to the seeding. Therefore, we set the rates of observation of metastases for such states to 0. We are interested in the distribution of the full system at the time of first observation, which can be triggered by either primary tumor or metastasis. We calculate this analogously to [34] as the stationary distribution 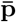 on the extended state space *𝒮*_*D*_ where each of the observation events halts the progression of the entire system. Each state where either observation occurred becomes an absorbing state. Thus the entire probability mass is located on the sets of states *O*_P_ = *𝒮 ×* (1, 0) (primary tumor is observed) and *O*_M_ = *𝒮 ×* (0, 1) (metastasis is observed). Analogous to Equation (5), we therefore have

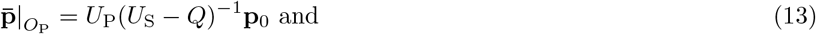

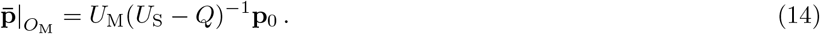

In most cases, there is a considerable time lag between the observation of a primary tumor and the observation of its metastatic offspring. To account for this, we model two consecutive observations. Consider the case where the primary tumor is observed first with genotype *x*^P^ *∈ {*0, 1*}*^*n*^ and the metastasis is only observed at a later point in time with genotype *x*^M^ *∈ {*0, 1*}*^*n*^. In this case the metastasis is unobservable at the time of primary tumor observation, and thus we are interested in the metastasis marginal probability 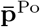 of only observing a primary tumor *x*^P^, given by

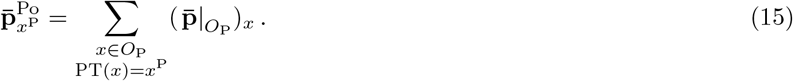

Note that each tumor in a dataset is observed exactly once and no information about its subsequent progression is available. Therefore we do not track the progression of the primary tumor after its observation. Instead from here on, we only model the progression of the still unobserved metastasis. To do so, we first calculate the distribution of metastasis genotypes at the time of primary tumor observation conditioned on the observed primary tumor genotype, which is given by

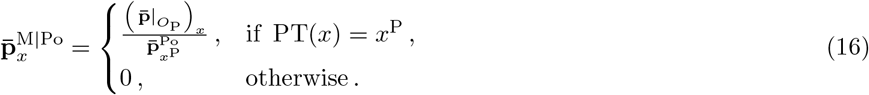

In words, we set the probability of all states where the primary tumor genotype is not equal to the observation to 0, and then renormalize the resulting vector to obtain the desired conditional distribution. Next analogously to [34] we propagate the distribution of unobserved metastases forward in time, until the metastasis is observed. This yields

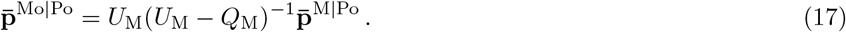

Finally, the probability to observe a primary tumor and metastasis pair in state *x*, given that the primary tumor was observed first is

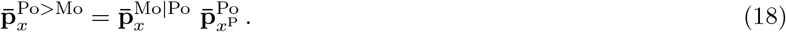

By an analogous calculation the probability to observe a primary tumor and metastasis pair in state *x*, given that the metastasis was observed first is given by

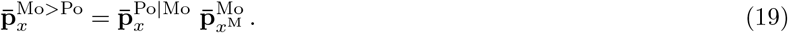

If the order of observation is not recorded then we evaluate the total probability to observe state *x* as

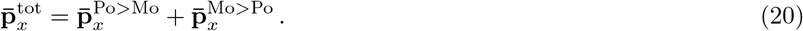

Equations (18), (19), (20) give the probabilities of observing pairs of genotypes. However, often only a single genotype is available, whereas the other is missing. Such individual data points are incorporated by first calculating the full joint distributions over all states and then by marginalizing over the missing genotypes. First consider the case, where only a primary tumor is observed with genotype *x*^P^, then marginalization over the unobserved metastasis genotypes yields

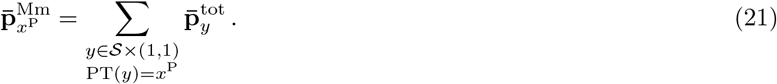

If a metastasis was observed but not sequenced, then we do not need to sum over all states, but only over states in *𝒮*_1_. Conversely, if evidence for the complete absence of metastases is available, then we only need to sum over states in *𝒮*_0_. Next, consider the case where only a metastasis is observed with genotype *x*^M^, then marginalizing over the unobserved primary tumor genotypes yields

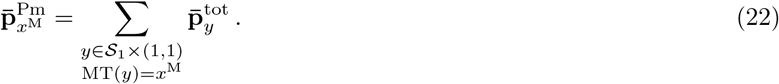

Since a metastasis is observed, we know that seeding must have occurred and therefore we only need to sum over states in *𝒮*_1_.

#### 2.2.3 Parameter estimation

The average log-likelihood of a dataset *𝒟* containing primary tumor and metastasis pairs as well as single genotypes is given by

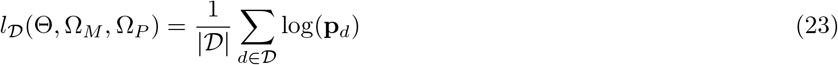

Where

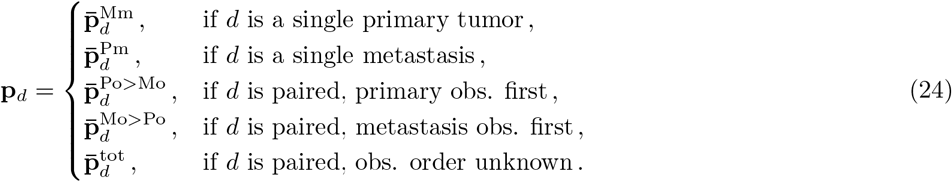

We infer the parameters Θ, Ω_M_, Ω_P_ from data via maximum likelihood estimation. We follow [34] and utilize the penalization

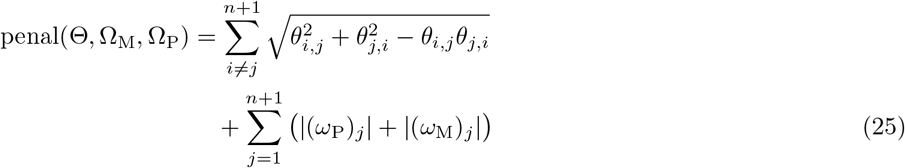

with *θ*_*i,j*_ = log(Θ_*i,j*_), (*ω*_M_)_*j*_ = log((Ω_M_)_*j*_), (*ω*_P_)_*j*_ = log((Ω_P_)_*j*_). The penalty promotes sparsity as the logarithmic parameters are shrunk to 0. Additionally, it promotes symmetry as effects between events *i* and *j* are grouped and selected together. We then optimize

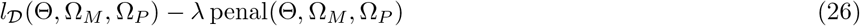

via gradient ascent. The hyper parameter *λ* is selected via 5-fold cross validation.

## 3 Results

To further our understanding of metastatic spread in lung adenocarcinomas, we trained metMHN on 4,852 paired and unpaired samples from the LUAD cohort of the MSK-IMPACT study. Next, we describe the dataset and then present our key findings.

### 3.1 Data preparation

We retrieved the AACR GENIE 14.1 data release [30] through synapse.org [12]. Our selection included all samples assayed at the Memorial Sloan-Kettering Cancer Center annotated with the ONCOTREE code ‘LUAD’ (Lung Adenocarcinoma). For primary tumors without corresponding metastasis samples, we retrieved information about their metastatic status from [26] and excluded samples where the status of the metastasis was unknown. The final dataset consisted of 453 matched primary tumor (PT)/metastasis (MT) samples, 2,127 unpaired MT samples, 595 PT samples without corresponding metastases (seeding=0), and 1,677 PT samples with metastases that were not sequenced (seeding=1). The three most highly mutated paired samples were excluded from our analysis due to computational challenges in processing them with metMHN. In total, our study included 2,725 PT and 2,580 MT samples from 4,852 patients. Metadata for each sample also included the age of the corresponding patient at which the sample was reported. This data informs the model about the order of observation of primary tumors and metastases in the same patients. When multiple PT or MT samples were present, we chose the PT sample with the youngest sampling age and the MT sample with the oldest sampling age.

Genomic data consisted of somatic mutation data and segmented log R ratio (LRR) copy number data derived from single-region bulk sequencing using the targeted MSK-IMPACT panel [11]. We annotated mutation data using OncoKB [8] and filtered for variants likely to be functional, as outlined in [34]. Our analysis was restricted to genes consistently included in all versions of the MSK-IMPACT panel [12]. Specifically, we examined mutations in the top 20 most frequently mutated genes. In the case of copy number alterations, we initially normalized segmented copy number data using mecan4CNA [15]. Amplifications were identified with LRR values corresponding to relative copy number gains *≥* 0.5. Conversely, deletions were marked by LRR values corresponding to relative copy number losses *≤ −*0.5. We determined the precise minimal intervals necessary for a copy number event classification in 8 instances, based on the minimal commonly altered regions per chromosome arm and gene extents. For amplifications, we required full gene extents to be covered by an alteration, whereas for deletions we allowed for shorter intervals. In total, our study considered 28 distinct genomic events, including mutational events (‘M’), copy number amplification (‘Amp’) and deletion (‘Del’) events. Binary event input data, alongside exact interval definitions for copy number events, records of the selected patients and samples and preparation scripts are accessible at https://github.com/cbg-ethz/metMHN.

### 3.2 Effects between genomic events and seeding

On the dataset described above, we fit metMHN and tuned the hyperparameter *λ* in a 5-fold cross-validation (Figure 2). Reassuringly, the LUAD model confirms several interactions well-documented in the literature. Specifically, it identifies the strong, antagonistic relationship (evidenced by a bidirectional negative edge) between the principal drivers KRAS (M) and EGFR (M) [37, 35]. Our model infers that EGFR suppresses further mutational co-drivers, which suggests that it might often be sufficient for progression. Instead, EGFR-driven LUADs frequently exhibit disruption of cell cycle regulation through copy number losses in RB1 and CDKN2A, two patterns also described in [25].

**Figure 2:**
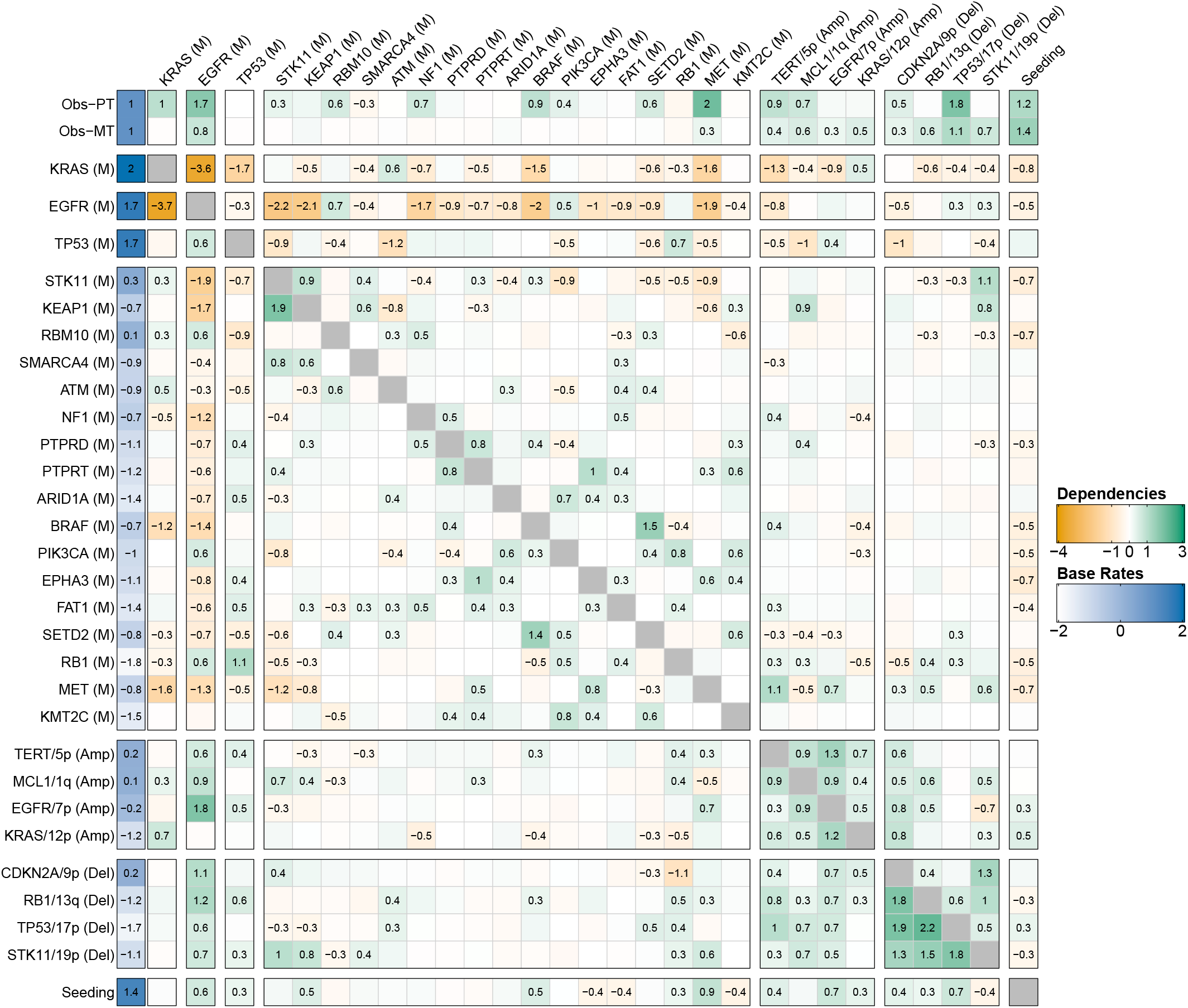
Interactions between progression events in lung adenocarcinomas. The log-effects on observation (clinical detection) of the primary tumor and metastasis *ω*_P_ and *ω*_M_ are plotted in the first two rows, the remaining matrix shows the log-interaction strengths among genomic events *θ*. The base rates of all events are plotted on the left (in blue). The effects an event *i* exerts on other events *j* are collected in the *i*^th^ column (outgoing edges). Vice versa the effects, that events *j* exert on event *i* are collected in the *i*^th^ row (incoming edges). Effects of genomic events on seeding are shown in the bottom row. Vice versa effects from seeding on genomic events are shown in the rightmost column.

The model further highlights synergistic interactions that reflect established oncogenic processes, such as the rate increases observed between STK11 (M) and KEAP1 (M), and between TP53 (M) and RB1 (M) [41, 6, 27]. metMHN also infers multiple positive interactions between gene mutations and corresponding copy number alterations, exemplified by the interaction between EGFR (M) and amplification of EGFR/7p, as well as between STK11 (M) and deletion of STK11/19p — a pattern commonly seen across various cancers [1]. Additionally, the model reflects that several mutational events capable of activating the (RTK)-RAS-RAF-MEK signaling pathway—namely, KRAS (M), EGFR (M), NF1 (M), BRAF (M), and MET (M)—tend to promote the observation of primary tumors and suppress each other’s occurrence [20].

### 3.3 metMHN identifies drivers of metastasis

We next examined the interactions between genomic events and metastatic seeding. The outgoing edges from the seeding event (rightmost column in Figure 2) represent the cancer cell’s adaptive response to the changing selective pressures encountered during its journey from the primary tumor to the metastatic site. The incoming edges into the seeding event (bottom row in Figure 2) indicate how particular mutations within the primary tumor may accelerate or impede the metastatic seeding rate, thereby pinpointing genetic elements that either drive or hinder metastasis development.

metMHN identifies mutations and amplifications in EGFR, along with TP53 mutations and deletions, and MET mutations, as accelerators of metastasis formation, as indicated by positive edges (i.e., promoting effects) from these events to the seeding event (Figure 2). These findings are substantiated by experimental evidence which indicate that activation of EGFR [36, 10], inactivation of TP53 [38, 29], and activation of MET [42, 9] enhance the metastatic capacity of lung cancer cells. Beyond these events, metMHN also revealed that various other copy number alterations positively influence the seeding process. Although widespread aneuploidy is typically regarded as a hallmark of advanced cancer stages [3], specific copy number changes, like CDKN2A deletions, have been documented to sometimes occur early in lung adenocarcinoma development [25, 39]. In this context we also note metMHN’s inference that copy number events generally do not substantially affect the primary tumor observation rate but indeed promote metastasis observability.

Interestingly, the effects promoting metastasis were relatively modest when compared to the base rate of seeding. This observation suggests that certain genetic or non-genetic drivers of the metastatic process might not be accounted for in the model. Alternatively, this could also indicate that primary tumor cells may inherently possess a propensity to metastasize, as suggested by [21]. Lastly, metMHN suggests that upon the seeding of metastases, the accumulation rates of many mutational events tend to decrease. This pattern could imply that once the metastatic process is initiated and in progress, there is diminished pressure for further mutational driver alterations, compared to the initial stages of primary tumorigenesis [13].

### 3.4 Relative timing of progression events and seeding

We computed the most likely chronological sequences of events for 313 paired data points and 2,127 unpaired metastases, where we limited our analysis to cases where calculations were feasible. For the paired data points the orderings are branched, as exemplified in Figure 3a. Prior to seeding, events happen jointly in the primary tumor. Upon seeding, the trajectory splits into a primary tumor branch and a metastasis branch (blue lower and red upper branches in Figure 3a, respectively). The unpaired metastases’ orderings are linear.

**Figure 3:**
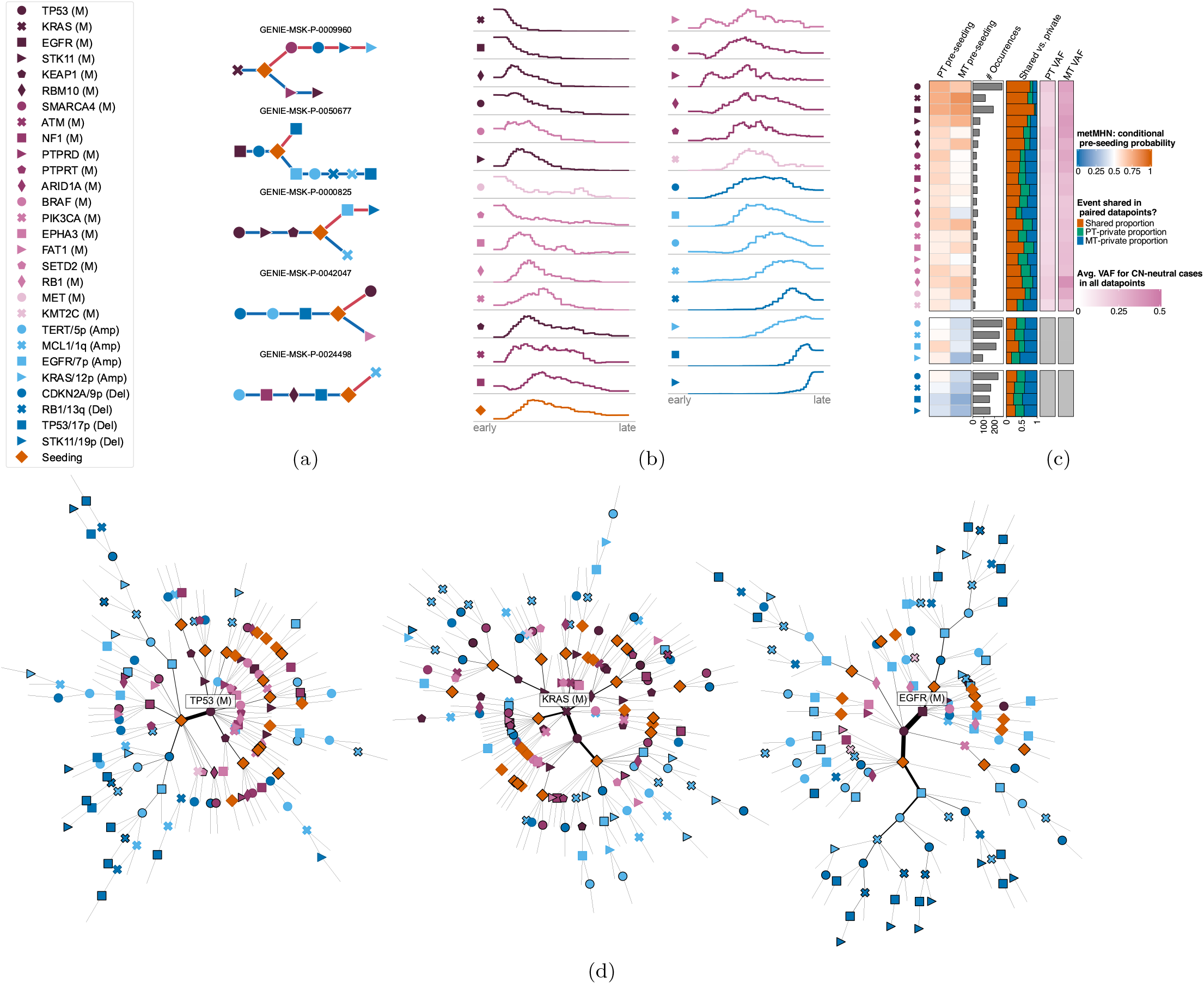
(a) Event orders for 5 patients as inferred by metMHN. Events accumulate from left to right. Blue edges represent the primary tumor development, red edges the one of the metastasis. Distances between events do not correspond to real or estimated time. (b) Distribution of relative positions in trajectories. The left end of the axes corresponds to the beginning, and the right to the end of progression. (c) Pre-seeding probabilities estimated by metMHN and empirical evidence from paired samples. The first and second column show the pre-seeding probabilities estimated by metMHN conditioned on the event being observed in the primary tumor (column 1) or the metastasis (column 2). Column 3 shows the number of occurrences for each event in the paired data, column 4 shows the proportions of shared versus private occurrences for each event in the paired data. Columns 5 and 6 show the mean variant allele frequencies in the primary tumor and the metastasis respectively. (d) Most probable event orderings for observed metastases genotypes as inferred by metMHN, stratified by TP53 (M) (left), KRAS (M) (middle) and EGFR (M) (right) as their first event. Each branch extending out from a tree’s root represents a group of metastases for which the events were inferred to occur in the order of the branch. Edge widths scale proportionally to the dataset’s count of metastases commencing with that particular sequence of events, and branches are trimmed at an edge threshold of 3. Black-bordered nodes indicate observed genotypes.

Next, we analyzed the distribution of event positions, relative to trajectory lengths (Figure 3b): The plots show for every event how often it occurred for each relative time point, where the left end of the axes corresponds to the beginning and the right to the end of progression. Well-established and highly frequent mutational drivers of LUAD progression, such as KRAS (M), EGFR (M) and TP53 (M) appear consistently early as initiating events. We find similar patterns for less frequent mutational events, such as MET (M) and SETD2 (M). Some events rarely appear as initiators, but still mostly occur in the early half of any sequence, such as STK11 (M) and BRAF (M). For example, RB1 (M) rarely happens spontaneously, which is reflected by its low base rate. However, it is promoted by both EGFR (M) and TP53 (M) and thus tends to happen subsequently, see Figure 2 and Figure 3d. Crucially, metastatic seeding was observed to happen at varying stages, with the majority of trajectories showing genomic progression both before and after seeding. On the late end of the spectrum we mainly find copy number events. After the first such event happens, it usually promotes other copy number events (see Figure 2), leading to compounding rate increases for copy number events towards the end of a typical trajectory, possibly reflecting genomic instability in late stage cancers [3].

Next, we stratified the inferred metastasis trajectories by the 3 most prevalent initial events. Specifically, trajectories starting with TP53 (M), KRAS (M) and EGFR (M) at the first position accounted for 1,766 patients or 72.38% of the analyzed metastases (Figure 3d). Remarkably, the subset of trajectories initiated by TP53 (M) (left side) included a significant number of tumors which seeded immediately after. These tumors then predominantly acquired copy number events. In a minority of cases, additional mutation events such as STK11 (M) and KEAP1 (M) occurred before seeding. Trajectories that began with KRAS (M) (center) generally showed later seeding, frequently after the accumulation of other mutational co-drivers, including TP53 (M), STK11 (M), KEAP1 (M), RBM10 (M), and ATM (M). These trajectories too typically concluded with a series of copy number events. Conversely, trajectories initiated by EGFR (M) (right side) exhibited distinctly different progression patterns. Contrary to those beginning with KRAS (M), these trajectories rarely accumulated additional mutational events before seeding, with TP53 (M) being an exception. Post-seeding, the progression was once again dominated by copy number changes. However, these events followed characteristic sequences, often starting with EGFR/7p (Amp) and CDKN2A/9p (Del), then proceeding to TP53/17p (Del) and STK11/19p (Del), and culminating with the clinical detection of the tumor.

### 3.5 metMHN is consistent with clonality information

A key quality of metMHN is its ability to quantify the timing of seeding relative to other progression events. To validate this, we compared it with an orthogonal readout of metastatic development relative to mutational events: A mutation that predates the seeding of a primary tumor clone is expected to be clonal, i.e., exhibit a high variant allele frequency (VAF, close to 0.5) in subsequent metastases [4]. In contrast, mutations arising post-seeding in metastases are more likely to be subclonal and thus exhibit lower VAFs. Therefore, we used per-gene mean VAFs in metastasis samples as a proxy for the relative timing (pre- or post-seeding) of the occurrence of mutations in the respective gene. To account for a bias in VAF distributions, we restricted VAF measurements to cases in which the respective gene was not copy number altered. We compared for each mutation its mean VAF with the model-derived probability that the event occurred prior to seeding. To this end, we approximated this probability through simulations using Gillespie’s algorithm [17]. We found that mutational events with high pre-seeding probabilities in metastases corresponded to elevated VAFs in metastasis samples as evidenced by a Pearson correlation coefficient of 0.55 (*p* = 0.01) see Figure 3c and Figure 1 in the supplement. In summary, while metMHN builds on co-occurrence patters and does not leverage VAF information, they nevertheless produce results consistent with clonality information.

## 4 Discussion

We have presented metMHN, an efficient analytical model for cancer progression, specifically designed to investigate the forking progression paths of primary tumors and their metastatic offspring. Distinguishing itself from specialized phylogenetic methods operating on rare multi-region sequencing data, metMHN capitalizes on the extensive cross-sectional data available from clinical targeted sequencing and is able to infer relationships between events that are shared across individual samples. Our comprehensive analysis, encompassing data from nearly 5000 lung cancer patients, corroborates well-established relationships among key genomic drivers. In addition, metMHN successfully identifies specific events in primary tumors that may accelerate the development of metastases and quantifies how the dynamics of event accumulation change upon metastatic branching. Moreover, metMHN allows for the reconstruction as well as for the simulation of disease histories yielding further insight into the dynamics of metastatic cancers. This dual capability of metMHN not only deepens our comprehension of the key events that propel cancer progression but also provides a quantitative perspective on how these interactions manifest into distinct histories of tumor progression.

Every model’s efficacy is inherently tied to the quality of its training data. While metMHN uses comprehensive cross-sectional data from bulk tissue, this approach has its limitations, particularly in resolving the clonal structures of heterogeneous tumors. In metMHN, binary states represent the tumor as a whole. Consequently, two tumors with identical mutations will be interpreted identically by the model, even if, in one case, the mutations exist within the same clone, and in the other, they are in separate clones. Another challenge arises when the training data does not accurately represent the patient population. For instance, an under-representation of metastatic tumors in the training data could lead to an underestimation of the base rate for the seeding event, falsely suggesting they occur later in the progression than they actually do, while an over-representation of these cases would have the opposite effect. In contrast, phylogenetic methods, which reconstruct tumor evolution on an individual basis, are less susceptible to biases in datasets. These methods also offer the advantage of resolving clonal structures, presenting a more detailed picture of tumor evolution. However, the scarcity of data, especially in multi-region sequencing studies, limits their ability to represent patient populations comprehensively.

In summary, metMHN complements phylogenetic analyses and stands out as the only cancer progression models capable of fully utilizing the largest clinical genomic datasets currently available. metMHN models offer a distinct advantage: They provide a quantitative and dynamic description of metastatic cancer progression. This unique approach enables them to contribute valuable insights into the complexities of metastatic spread, enriching our understanding of cancer progression with their analytical perspective.

## 5 Competing interests

No competing interest is declared.

## 6 Author contributions statement

KR, RSC, NB and RSP conceptualized and initiated the project. KR, RSC and YLH developed the model. KR, YLH, CN implemented the algorithms. SP, MK, TW and LG provided numerical foundations for model analysis. AL prepared the input data. AL, YLH, KR analysed the LUAD data. KR, AL, YLH, and RSP drafted the manuscript. All authors critically read and improved upon the draft.

## 7 Acknowledgments

We thank Nikolaus Schultz from MSKCC, Thomas Ratzke, Stefan Vocht and Stefan Hansch for helpful discussions. This work is supported by the Deutsche Forschungsgemeinschaft (DFG: TRR 305-A02), GR-3179/6-1 “Tensorapproximation-smethoden zur Modellierung von Tumorprogression” Swiss National Science Foundation grant 179518 and Swiss Cancer League grant KFS-2977-08-2012.

## Notes

### Competing Interest Statement

The authors have declared no competing interest.

