## Supplemental Files for "Modeling metastatic progression from cross-sectional cancer genomics data"

### 1 Sum of tensor product formulations

Here we provide formulas for the transition rate matrices $Q_0, Q_\mathrm{P}, Q_\mathrm{M}, Q_\mathrm{S}$ as sums of tensor products. We use "$\cdot$" to denote 0.

$$Q_0 = \sum_{i=1}^{n} \left( \bigotimes_{j=1}^{i} \begin{pmatrix} 1 & \cdot & \cdot & \cdot \\ \cdot & \cdot & \cdot & \cdot \\ \cdot & \cdot & \cdot & \cdot \\ \cdot & \cdot & \cdot & \Theta_{i,j} \end{pmatrix} \otimes \begin{pmatrix} -\Theta_{i,i} & \cdot & \cdot & \cdot \\ \cdot & \cdot & \cdot & \cdot \\ \cdot & \cdot & \cdot & \cdot \\ \Theta_{i,i} & \cdot & \cdot & \cdot \end{pmatrix} \otimes \bigotimes_{j=i+1}^{n} \begin{pmatrix} 1 & \cdot & \cdot & \cdot \\ \cdot & \cdot & \cdot & \cdot \\ \cdot & \cdot & \cdot & \cdot \\ \cdot & \cdot & \cdot & \Theta_{i,j} \end{pmatrix} \right) \otimes \begin{pmatrix} 1 & \cdot \\ \cdot & \cdot \end{pmatrix} \tag{1}$$

$$Q_\mathrm{S} = \Theta_{n+1,n+1} \bigotimes_{j=1}^{n} \begin{pmatrix} 1 & \cdot & \cdot & \cdot \\ \cdot & \cdot & \cdot & \cdot \\ \cdot & \cdot & \cdot & \cdot \\ \cdot & \cdot & \cdot & \Theta_{n+1,j} \end{pmatrix} \otimes \begin{pmatrix} -1 & \cdot \\ 1 & \cdot \end{pmatrix} \tag{2}$$

$$Q_\mathrm{P} = \sum_{i=1}^{n} \left( \bigotimes_{j=1}^{i} \begin{pmatrix} 1 & \cdot & \cdot & \cdot \\ \cdot & \Theta_{i,j} & \cdot & \cdot \\ \cdot & \cdot & 1 & \cdot \\ \cdot & \cdot & \cdot & \Theta_{i,j} \end{pmatrix} \otimes \begin{pmatrix} -\Theta_{i,i} & \cdot & \cdot & \cdot \\ \Theta_{i,i} & \cdot & \cdot & \cdot \\ \cdot & \cdot & -\Theta_{i,j} & \cdot \\ \cdot & \cdot & \Theta_{i,j} & \cdot \end{pmatrix} \otimes \bigotimes_{j=i+1}^{n} \begin{pmatrix} 1 & \cdot & \cdot & \cdot \\ \cdot & \Theta_{i,j} & \cdot & \cdot \\ \cdot & \cdot & 1 & \cdot \\ \cdot & \cdot & \cdot & \Theta_{i,j} \end{pmatrix} \right) \otimes \begin{pmatrix} \cdot & \cdot \\ \cdot & 1 \end{pmatrix} \tag{3}$$

$$Q_\mathrm{M} = \sum_{i=1}^{n} \left( \bigotimes_{j=1}^{i} \begin{pmatrix} 1 & \cdot & \cdot & \cdot \\ \cdot & 1 & \cdot & \cdot \\ \cdot & \cdot & \Theta_{i,j} & \cdot \\ \cdot & \cdot & \cdot & \Theta_{i,j} \end{pmatrix} \otimes \begin{pmatrix} -\Theta_{i,i} & \cdot & \cdot & \cdot \\ \cdot & -\Theta_{i,j} & \cdot & \cdot \\ \Theta_{i,j} & \cdot & \cdot & \cdot \\ \cdot & \Theta_{i,j} & \cdot & \cdot \end{pmatrix} \otimes \bigotimes_{j=i+1}^{n} \begin{pmatrix} 1 & \cdot & \cdot & \cdot \\ \cdot & 1 & \cdot & \cdot \\ \cdot & \cdot & \Theta_{i,j} & \cdot \\ \cdot & \cdot & \cdot & \Theta_{i,j} \end{pmatrix} \otimes \begin{pmatrix} \cdot & \cdot \\ \cdot & \Theta_{i,n+1} \end{pmatrix} \right) \tag{4}$$

Here we additionally provide formulas for the diagonal matrices $U_\mathrm{P}$ and $U_\mathrm{M}$, that hold the observation rates, as sums of tensor products:

$$U_\mathrm{P} := \bigotimes_{i=1}^{n} \begin{pmatrix} 1 & \cdot & \cdot & \cdot \\ \cdot & \Omega_{\mathrm{P}\,i} & \cdot & \cdot \\ \cdot & \cdot & 1 & \cdot \\ \cdot & \cdot & \cdot & \Omega_{\mathrm{P}\,i} \end{pmatrix} \otimes \begin{pmatrix} \Omega_{\mathrm{P}\,n+1} & \cdot \\ \cdot & \Omega_{\mathrm{P}\,n+1} \end{pmatrix} \tag{5}$$

$$U_\mathrm{M} := \bigotimes_{i=1}^{n} \begin{pmatrix} 1 & \cdot & \cdot & \cdot \\ \cdot & 1 & \cdot & \cdot \\ \cdot & \cdot & \Omega_{\mathrm{M}\,i} & \cdot \\ \cdot & \cdot & \cdot & \Omega_{\mathrm{M}\,i} \end{pmatrix} \otimes \begin{pmatrix} \cdot & \cdot \\ \cdot & \Omega_{\mathrm{M}\,n+1} \end{pmatrix} \tag{6}$$

### 2 Supplementary plots

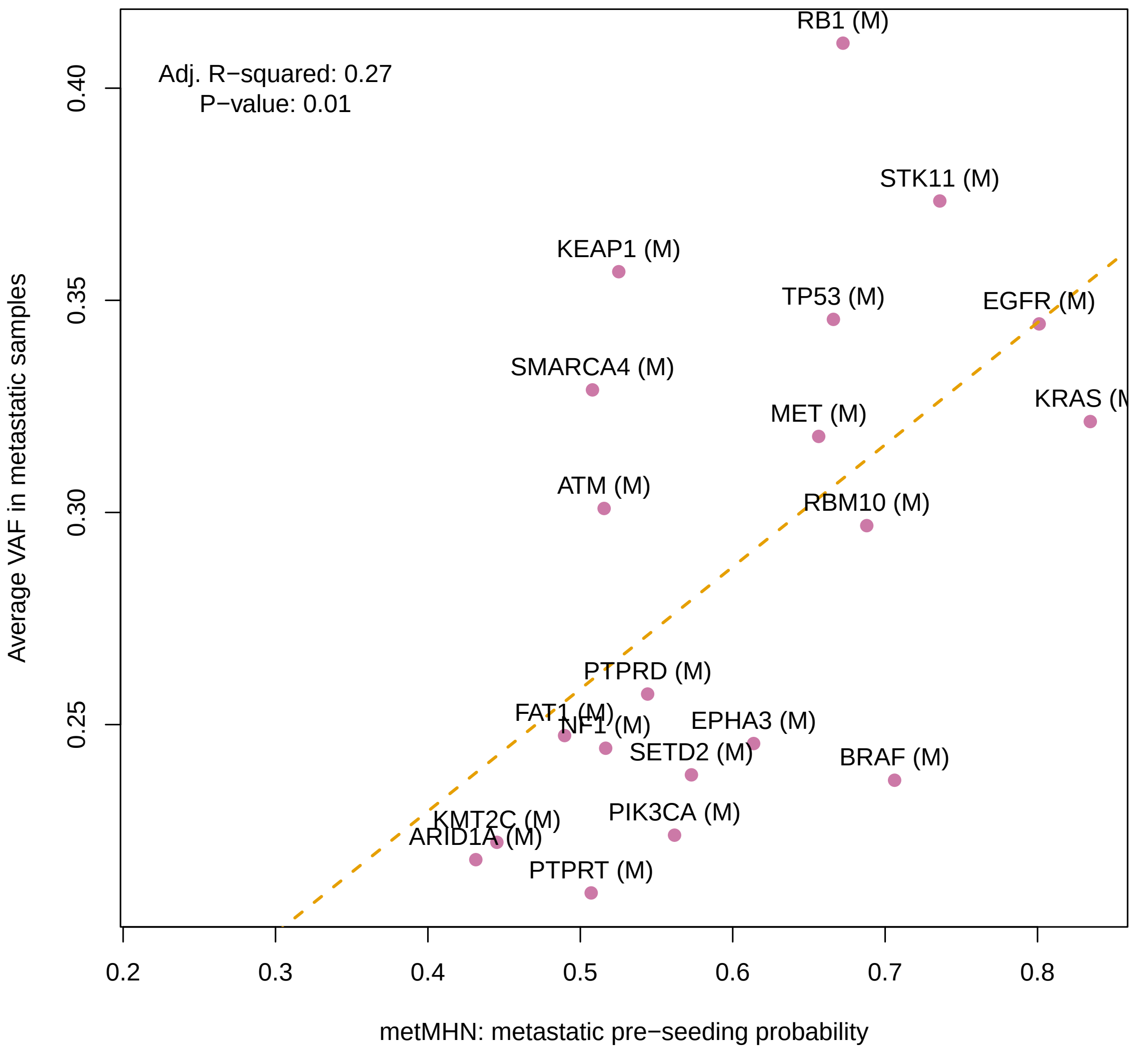


Figure 1: Metastasis pre-seeding probabilities for each mutational event, plotted against the average variant allele frequencies in copy number neutral cases for the respective event. A linear regression model was fit to evaluate the association, with an adjusted R-squared value of 0.27 and a p-value of 0.01.
